# Super-resolution analyzing spatial organization of lysosomes with an organic fluorescent probe

**DOI:** 10.1101/2022.02.14.480395

**Authors:** Lei Wang, Rui Chen, Guanqun Han, Xuan Liu, Taosheng Huang, Jiajie Diao, Yujie Sun

**Author notes:** Corresponding authors: Yujie Sun or Jiajie Diao. These authors contribute equally (Lei Wang, Rui Chen).

## Abstract

Lysosomes are multifunctional organelles involved in macromolecule degradation, nutrient sensing and autophagy. Live imaging has revealed lysosome subpopulations with dynamics and characteristic cellular localization. An as-yet unanswered question is whether lysosomes are spatially organized to coordinate and integrate their functions. Combined with super-resolution microscopy, we designed a small organic fluorescent probe, TPAE, that targeted lysosomes with a large Stokes shift. When we analyzed the spatial organization of lysosomes against mitochondria in different cell lines with this probe, we discovered different distance distribution patterns between lysosomes and mitochondria during increased autophagy flux. By using *SLC25A46* mutation fibroblasts derived from patients containing highly fused mitochondria with low oxidative phosphorylation, we concluded that unhealthy mitochondria redistributed the subcellular localization of lysosomes, which implies a strong connection between mitochondria and lysosomes.

## 1. Introduction

Lysosome is a well-known membrane-enclosed cytoplasmic organelle with more than 60 acid hydrolases in its lumen that is responsible for the degradation of all kinds of biological macromolecules.^[1]^ Due to its role in terminal degradation, lysosome is often referred as a cell’s “garbage disposal” system.^[1]^ In the past, lysosome was only considered to be a static organelle dedicated to recycling wastes in cells, but new discoveries have revealed that lysosomes are dynamic structures that participate in many other cellular processes, such as cell metabolism mediation, signaling cell pathways, and transcription, to maintain organismal homeostasis. Lysosomes form membrane contact sites to exchange content and information with other cellular organelles, such as mitochondria. Mitochondria are necessary for cellular respiration and also function as storage for metabolites such as calcium, iron, lipids, protons, and ATP.^[2]^

Several recent studies have utilized different imaging techniques to observe the contacts formed between mitochondria and lysosomes;^[3]^ it was determined that the mitochondria-lysosome contacts (MLCs) are mediated by multiple proteins on both the mitochondrial and lysosomal membranes. To date, it has been known that the switch between Rab7 GTP-binding and GDP-binding states is a master regulator for the MLCs.^[4]^ And MLCs are very critical for mitochondrial functions, like fission^[4-5]^ and mitochondrial Ca^2+^ dynamics.^[6]^ However, the spatial organization between lysosomes and mitochondria is not clear.

Super-resolution microscopy bypasses the diffraction limit in light microscopy and enables the visualization of previously invisible molecular details in biological systems.^[7]^ Among the super-resolution techniques, structured illumination microscopy (SIM) is popular for researchers because of its many advantages, such as wavelength diversity, fast imaging, and simple sample preparation. Due to its better resolution, super resolution imaging has been used extensively to investigate cellular contents.^[8]^ Especially, the viscosity of mitochondria and lysosomes in live cells has been reported.^[9]^ However, few studies have been dedicated to reveal the spatial organization between lysosomes and mitochondria using SIM techniques.

In this study, we designed a small organic fluorescent probe, TPAE, which possesses excellent photophysical properties for SIM imaging. Due to its large Stokes shift, TPAE can be employed in super-resolution imaging when there is extremely low background fluorescence. Importantly, we were able to utilize TPAE to analyze the behavior of lysosomes in different biological processes and revealed a specific organization of lysosomes during autophagy.

## 2. Results and Discussion

### 2.1 TPAE Synthesis and Characterization

The TPAE synthesis started from commercially available triphenylamine; the intermediate 4,4’-((4-bromophenyl)azanediyl)dibenzaldehyde was obtained following a two-step reported procedure (Figure S1), which was subjected to a Suzuki coupling with pyridine-4-ylboronic acid afforded TPACHO.^[10]^ In the presence of malonic acid, the aldehyde groups in TPACHO were further converted to the cinnamic acid groups due to a Knoevenagel condensation reaction to form TPACOOH with a 60% yield. Finally, an esterification reaction between methanol and TPACOOH furnished the designed probe TPAE with a yield of 42% (Figure 1A). The physical characterization results of the synthetic intermediates and TPAE are included in Figures S2-S7.

**Figure 1.**
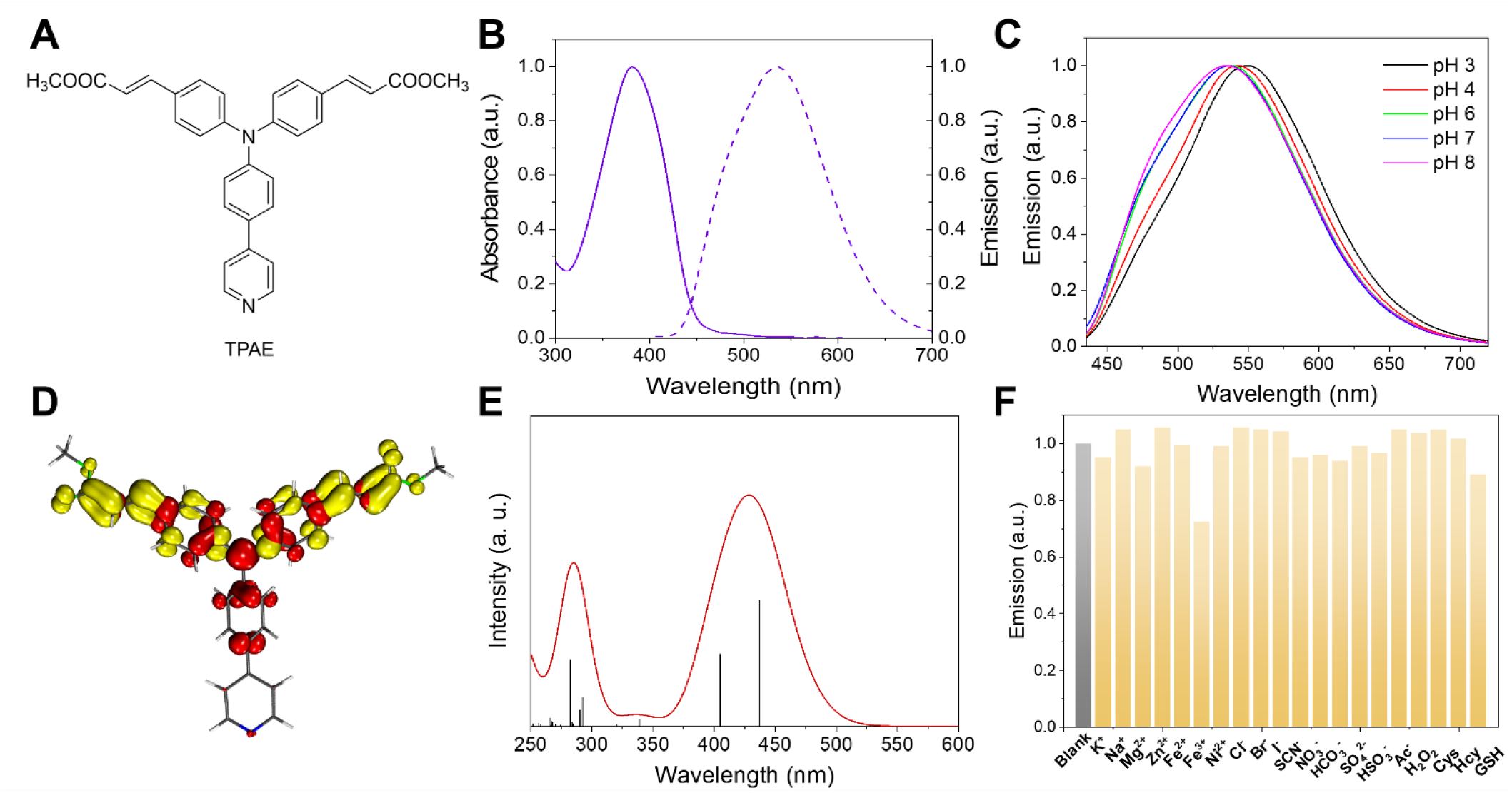
Structure and characterization of TPAE. (A) The chemical structure of TPAE. (B) Absorption (solid line) and emission (dash line, λ_ex._= 380 nm) spectra of TPAE in aqueous buffer at Ph 7. (C) Emission spectra of TPAE in aqueous buffers of different pH levels. (D) EDDM of the first singlet excited state (S_1_) of TPAE: red and yellow indicate decrease and increase in electron density, respectively. (E) Calculated absorption spectrum of TPAE (solvent = water, isovalue = 0.04). (F) Emission intensity of 10 μM TPAE probe in water after the addition of various chemical species. Conditions: the concentrations of agents from K^+^ to H_2_O_2_ are 100 μM, and from Cys to GSH are 200 μM, λ_ex_ = 380 nm, λ_em_ = 541 nm.

The UV-vis absorption spectrum of TPAE measured in an aqueous pH 7 buffer revealed an absorption peak at 380 nm (Figure 1B) with a molar extinction coefficient of 42,590 M^-1^cm^-1^ (Figure S8). Upon excitation at 380 nm, a strong emission was obtained and its emission maximum occurred at 540 nm. Hence, TPAE demonstrated a large Stokes shift of 160 nm, which rendered it a promising fluorescent probe in bioimaging applications with negligible background fluorescence.^[11]^ Moreover, the emission quantum yield of TPAE was measured at 12.5% using the commercial dye Coumarin 153 as a reference (ϕ = 12%).^[12]^

Because photostability is critically important for the application of organic fluorescent probes in biological media, we first measured the absorption and emission spectra of TPAE in aqueous buffers of different pH levels. As shown in Figure S9, no apparent changes were observed for the absorption spectra between pH 3–8, and the emission profile only exhibited a negligible shift in the same pH region (Figure 1C). Overall, these results suggest that an acidic environment does not strongly affect the photophysical properties of TPAE in aqueous media. To shed light on the pH-independent photophysical properties of TPAE, we performed density functional theory computations for TPAE. As delineated in Table S1, the highest occupied molecular orbital (HOMO) of TPAE is primarily located in the central triphenylamine moiety and the lowest unoccupied molecular orbital (LUMO) is primarily populated on the two side ester moieties. It should be noted that the terminal pyridine group was not involved in either the HOMO or the LUMO. The lowest singlet excited state (S_1_) of TPAE was computed to be largely composed of the electronic transition from the HOMO to the LUMO (99%) and located at 437 nm (Table S2). The calculated electron density difference map (EDDM) of the S_1_ state vividly presents the electron density change during this electronic transition (Figure 2D), where red and yellow indicate decrease and increase in electron density, respectively. The results of the simulated absorption spectrum of TPAE based on theoretical computations are plotted in Figure 2E. Since the terminal pyridine unit of TPAE is not involved in the S_1_ state, it is unsurprising that the absorption and emission of TPAE exhibited negligible pH dependence.

**Figure 2.**
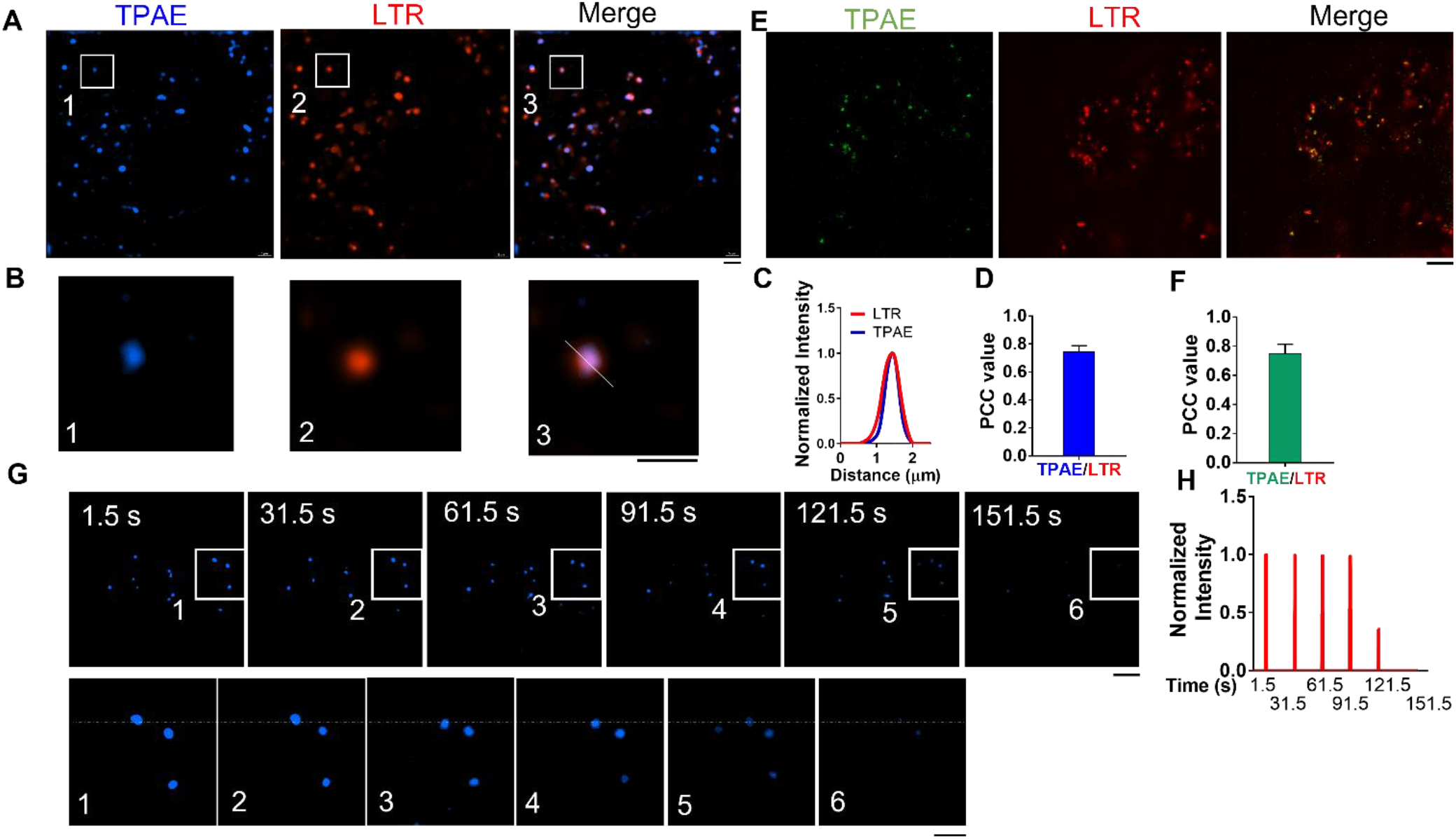
SIM characterization of TPAE in living cells. (A) Super resolution imagining of HeLa cells stained with TPAE (405 nm laser excitation, 420-495 nm emission) and LysoTracker-red (LTR) (561 nm laser excitation, 570-640 nm emission). Scale bar, 5 μm. (B) Zoomed-in images of white rectangles in A) Scale bar, 3 μm. (C) Fluorescence intensity profiles in white line from (B). (D) Pearson correlation coefficient (PCC) value for TPAE (405 nm laser excitation, 420-495 nm emission) and LTR in HeLa cells. n = 6, and data are presented as mean ± SEM. (E) Super resolution imagining of HeLa cells stained with TPAE (405 nm laser excitation, 500-550 nm emission) and LTR (561-nm laser with band path 570-640 nm). Scale bar, 5 μm. (F) Pearson correlation coefficient (PCC) value for TPAE (561 nm laser excitation, 570-640 nm emission) and LTR in HeLa cells. n = 6, and data are presented as mean ± SEM. (G) Photobleaching properties of TPAE for imaging under 405 nm laser excitation. Scale bars, 5 μm (upper panel) and 3 μm (lower panel). (H) Fluorescence intensity plot from the white dotted line in (F).

Next, we investigated the impact of potential interfering species on TPAE’s emission intensity in water. As displayed in Figure 1F, various biologically-related chemical species, including metal cations (K^+^, Na^+^, Mg^2+^, Zn^2+^, Fe^2+^, Fe^3+^, Ni^2+^), anions (Cl^-^, Br^-^, I^-^, SCN^-^, NO_3_^-^, HCO_3_^-^, SO_4_^2-^, HSO_3_^-^, Ac^-^), reactive oxygen species (H_2_O_2_), and reactive sulfur species (Cys, Hcy, GSH) were added to the TPAE solution; none of these interfering species were able to quench the strong TPAE emission in water upon excitation at 380 nm. Besides, we collected the absorption and emission spectra of TPAE along the increasing addition of glycerol in water. As can be observed in Figure S10, only a very slight red shift was observed for the absorption spectra (Figure S10A); in contrast, no linear change was observed for the emission spectra (Figure S10B). Regardless, strong emission intensity was retained with similar emission maxima (540-550 nm). Finally, the photostability test of TPAE was investigated under an irradiation wavelength of 405 nm. As depicted in Figure S11, emission intensity was almost unchanged after 300 s.

### 2.2 TPAE SIM Characterization in Living Cells

To attain a proper amount of TPAE to practice super-resolution imaging on cells, we examined the cytotoxicity of TPAE on HeLa cells. There was no difference on the viability of HeLa cells after being treated with TPAE within the concentration range of 0-100 μM after incubating for 24 h (Figure S12). To characterize the imaging characteristics of TPAE in living cells, HeLa cells were stained with TAPE for 30 min at a concentration of 5 μM, after which they were directly observed under structure illumination microscopy (SIM). The results revealed that TPAE easily passes through the cell membrane and accumulates in organelles, most likely lysosomes (405 nm laser excitation, 420-495 nm emission (blue color), Figure 2A). To further confirm that the organelles targeted by TPAE are lysosomes, co-staining of TPAE with the commercial dye LysoTracker Red (LTR) was performed. We observed that TPAE co-localized well with LTR (Figures 2A and 2B) and obtained a slightly higher resolution than LTR (Figure 2C) with a Pearson correlation coefficient (PCC) value of ∼0.79 (Figure 2D). Given TPAE’s large Stokes shift (160 nm), it can effectively avoid the background noise caused by excited light. To illuminate this advantage, we chose 405 nm laser excitation with 500-550 nm emission (green color) to visualize TPAE with SIM. TPAE again co-localized well with LTR (Figure 2E) and reached a PCC value of ∼0.75 (Figure 2F), which was similar to the results obtained at the previous imaging setting of 405 nm laser excitation and 420-495 nm emission (Figure 2D). To characterize the photobleaching resistance of TPAE, we exposed them to SIM laser illumination and reached a lifetime of more than 120 s (Figures 2G-2H).

### 2.3 Spatial Organization of Lysosomes and Mitochondria in Healthy and Mitophagy Conditions

We then applied this probe in biological studies. Lysosomes have been reported to play an important role in the autophagy process.^[13]^ To investigate the behavior of lysosomes during this process, we induced mitophagy, a form of autophagy that selectively degrades damaged mitochondria, by damaging mitochondria with carbonyl cyanide *m*-chlorophenyl hydrazone (CCCP), a common mitophagy inducer.^[8b, 14]^ After treating HeLa cells with CCCP for 12 h, we observed that the shape of the mitochondria stained with MitoTracker Deep Red (MTDR) significantly changed and became shorter and more rounded (Figure 3A), and damaged mitochondria also lost their cristae in the inner membrane (Figures 3B and 3C).

**Figure 3.**
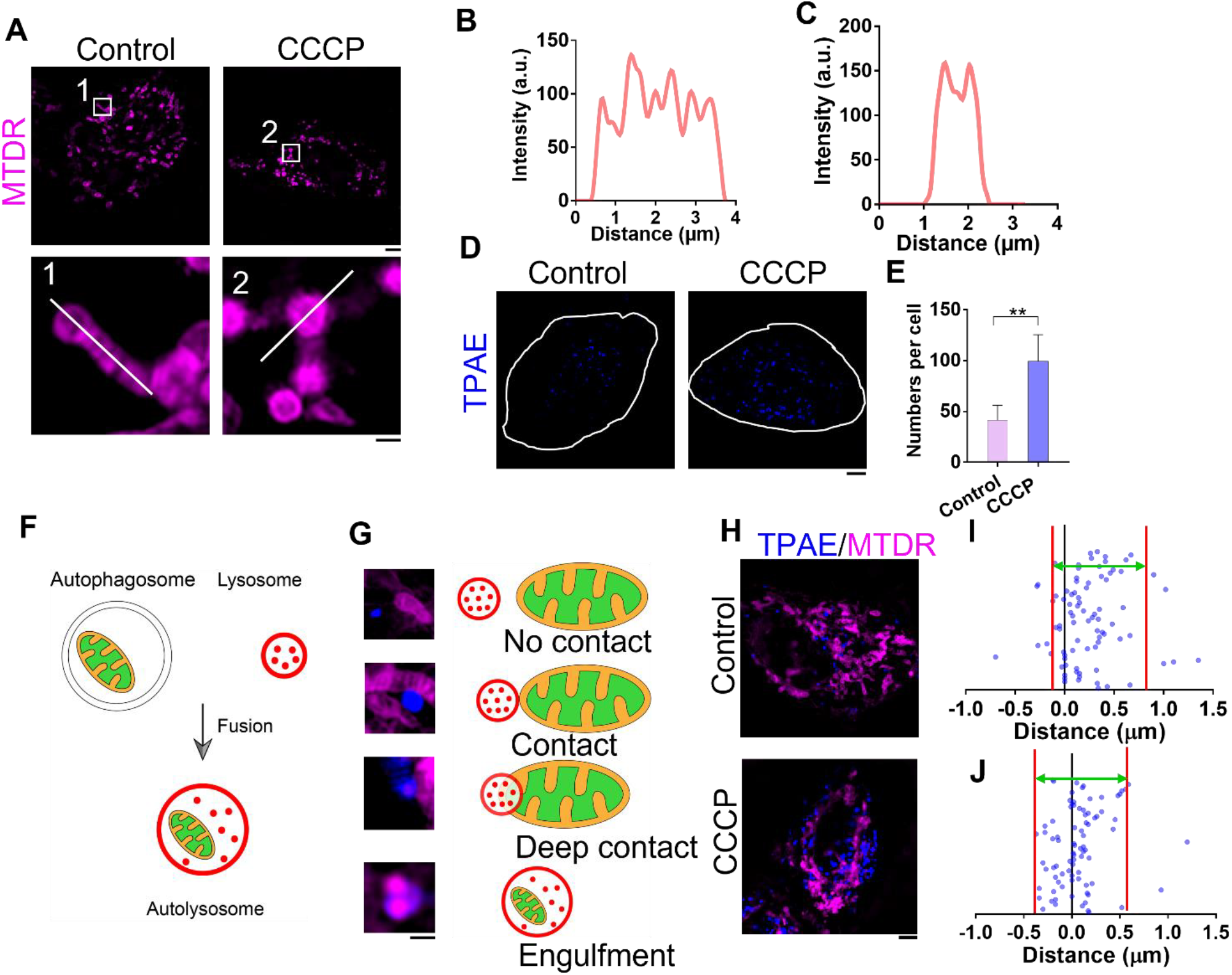
Analysis of interactions between TPAE-stained lysosomes and mitochondria in HeLa cells. (A) SIM images of HeLa cells stained with MitoTracker deep red (MTDR) after 12-hour treatment with or without CCCP. Scale bars, 5 μm (top panel) and 1 μm (lower panel). (B) Fluorescent plot from white line at No. 1 region of interest in (A). (C) Fluorescent plot from white line at No. 2 region of interest in (A). (D) SIM images of HeLa cells stained with TPAE (λ_ex_ = 405 nm, λ_em_ = 450 nm) after 12-hour treatment with or without CCCP, and white line indicates the outline of the cell. Scale bar, 5 μm. (E) Quantification of the number of lysosomes in HeLa with and without CCCP treatment; n = 10 cells, and data are presented as mean ± SEM; \*\**p <* 0.01, unpaired two-tailed t-test. (F) Schematic of mitochondria engulfed by autolysosome. (G) Types of mitochondria and lysosomes contact. Scale bar, 1 μm. (H) Representative SIM images of HeLa cells co-stained with TPAE and MTDR with or without 12-hour CCCP treatment. Scale bars, 5 μm. (I) and (J) Quantification of distance between mitochondria and lysosomes in HeLa cells without (I) or with CCCP treatment (J) (n = 84 events from eight cells); space between red lines indicates area where a majority of distances are distributed.

Even though the mitochondria undergo significant damages, the cells appear normal in bright field images (Figure S13). When we further examined the TPAE-stained lysosomes in the condition of mitophagy, we observed that the number of lysosomes in HeLa cells undergoing mitophagy dramatically increased (Figures 3D and 3E). When the mitophagy process was initiated, double-membraned autophagosomes enclosed the damaged mitochondria and fused them with lysosomes to form autolysosomes for degradation (Figure 3F). Due to the existence of autolysosomes, four types of physical interactions between mitochondria and lysosomes were observed: no contact, contact, deep contact, and engulfment (Figures 3G and 3H). We also discovered that the co-localization of mitochondria and lysosomes increased during mitophagy (Figure S14). Finally, the distance between mitochondria and their nearest lysosomes were quantified; the distances were shorter after the mitophagy process was induced in HeLa cells (Figures 3I and 3J), which displayed a unique pattern of spatial distribution of lysosomes in cells. This suggests that lysosomes adopt a different spatial organization during an increased level of autophagy.

### 2.4 Spatial Organization of Lysosomes and Mitochondria in Fibroblasts

We examined the lysosome behavior in wild type fibroblasts to determine whether lysosomes adopt different distribution patterns during mitophagy. Similar to the HeLa cell, mitochondria suffered dramatic changes in shape after the CCCP-induced damage (Figure 4A), yet these changes seemed to have no effect on the cell shape (Figure S15). Surprisingly, the number of lysosomes in fibroblasts significantly decreased after the CCCP treatment (Figure 4B), which suggests a regulation mechanism of lysosomes in fibroblasts that differs from the regulation mechanism in cancer-cell lines (Figure 3E). The distance between mitochondria and lysosomes became shorter during mitophagy in fibroblasts, which also happened in HeLa cells (Figures 4C, 4D and 4E). Notably, the co-localization of mitochondria and lysosomes further increased after the CCCP treatment (Figure S16).

**Figure 4.**
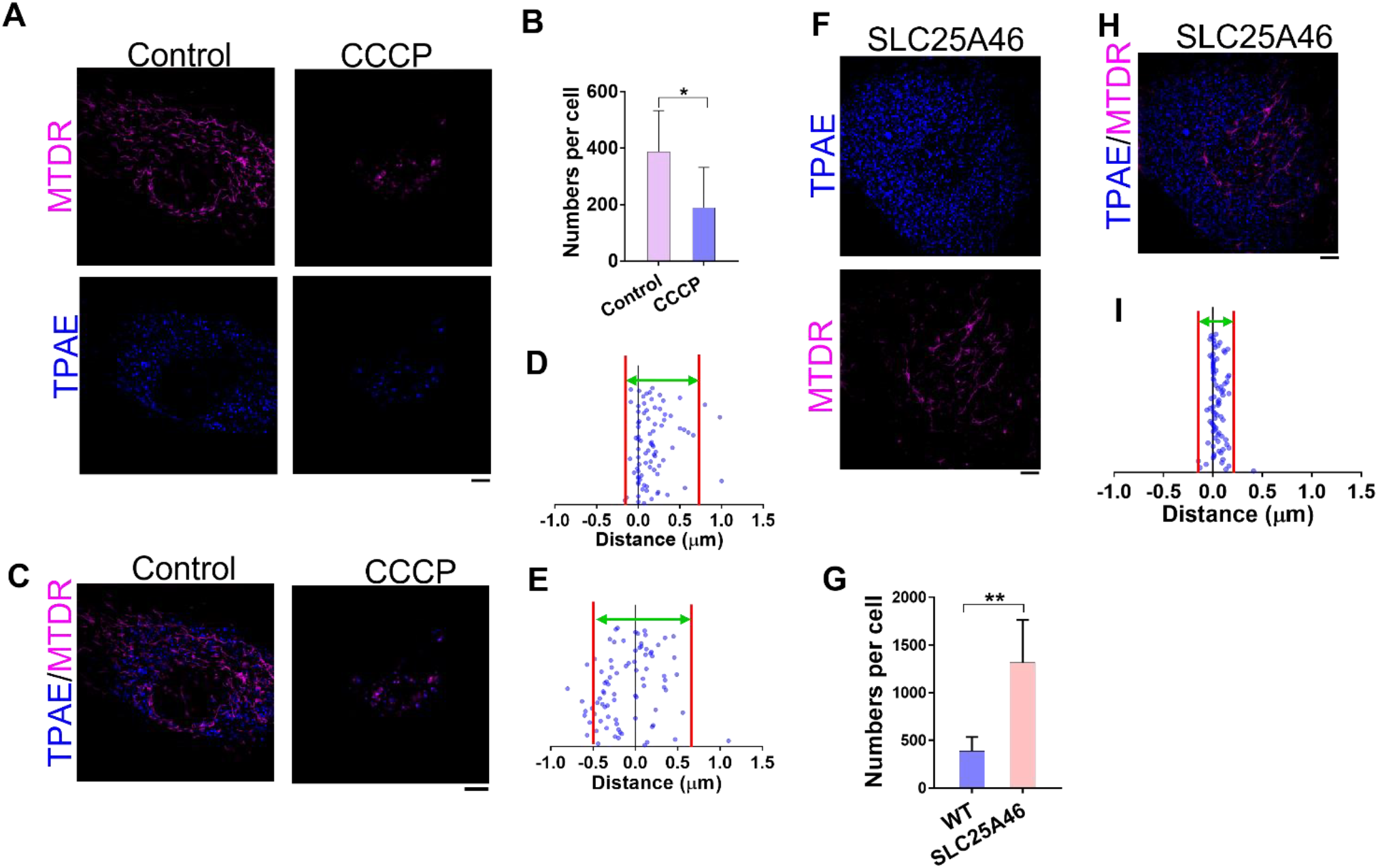
Analysis of mitochondria and lysosomes interactions in wild type (WT) human fibroblasts and patient-derived *SLC25A46* mutant fibroblasts. (A) WT fibroblasts stained with TPAE and MTDR, respectively, after a 12-hour treatment with or without CCCP. (B) Quantification of the number of lysosomes in WT fibroblasts with or without CCCP treatment; n = 10 cells, and data are presented as mean ± SEM; \**p <* 0.05, unpaired two-tailed t-test. (C) Representative images of co-staining of TPAE and MTDR with fibroblasts with or without CCCP treatment. (D) Quantification of distance between mitochondria and lysosomes in WT fibroblast cells without CCCP treatment (n = 84 events from five cells). (E) Quantification of distance between mitochondria and lysosomes in WT fibroblast cells with CCCP treatment (n = 84 events from five cells). (F) Patient-derived *SLC25A46* mutant fibroblasts were stained with TPAE and MTDR, respectively. (G) Quantification of the number of lysosomes in WT and *SLC25A46* mutant fibroblasts, respectively; n = 10 cells, and data are presented as mean ± SEM; \*\**p <* 0.01, unpaired two-tailed t-test. (H) Representative images of TPAE and MTDR co-stained with *SLC25A46* mutant fibroblasts. (I) Quantification of distance between mitochondria and lysosomes in *SLC25A46* mutant fibroblasts (n = 84 events from five cells); space between the red lines indicates the area where a majority of distances in (D), (E), and (I) are distribute. Scale bars, 5 μm.

Due to the strong connection between mitochondria and lysosomes, we exploited patient-derived *SLC25A46* mutant fibroblasts; this led to highly fused mitochondria and decreased mitochondrial oxidative phosphorylation and resulted in clinical symptoms of mitochondrial diseases.^[8g]^ This gene encodes a mitochondrial solute carrier protein family member, and its mutation results in neuropathy and optic atrophy. Even though the cell shapes of the *SLC25A46* mutant fibroblasts were not different from the wild type (WT) (Figure S17), the mitochondria were relatively longer than those in the WT fibroblasts. There were more lysosomes in mutants than in the WT (Figure 4G), however, which implies that unhealthy mitochondria strongly influence lysosomes. We again measured the distance between the mitochondria and the nearest lysosomes and obtained a unique lysosome distribution pattern in the *SLC25A46* mutant fibroblasts (Figures 4H and 4I), compared to what was found in the WT (Figure 4D), which suggests that the mitochondria had a closer contact with the lysosomes in mutant cell lines. Interestingly, the co-localization of lysosomes and mitochondria were retained in a similar fashion by both the WT and the mutants (Figure S18). Taken together, our results strongly indicate that mitochondria are able to regulate the spatial distribution of lysosomes.

## 3. Conclusion

To summarize, we designed a small fluorescent probe TPAE with a large Stokes shift that could be readily used for super resolution microscopy; the photophysical properties of this probe were well-characterized and -investigated by utilizing this probe in biological studies. When the behavior of lysosomes was analyzed during the autophagy process, a unique spatial organization pattern of lysosomes against mitochondria was discovered. Our results revealed that the biogenesis rate of lysosomes in fibroblasts during autophagy is different from that in cancer cells, and the dynamics of lysosomes are not a random event; instead, they move closer to the mitochondria when the mitochondria are damaged. Finally, we found that endogenously unhealthy mitochondria affect the distribution of lysosomes, which implies a new regulation mechanism between lysosomes and mitochondria that awaits further investigation.

## Acknowledgements

Y.S. acknowledges the University of Cincinnati and the partial support of the National Science Foundation (CHE 1955358). J.D. thanks the National Institute of Heath (R35GM128837). The OMX-SR 3D-SIM microscope at the Light Microscopy Imaging Center, Indiana University was provided by the NIH grant NIH1S10OD024988-01.

