## Supplementary material for "Super-resolution analyzing spatial organization of lysosomes with an organic fluorescent probe": SI Figures

### These authors contribute equally.

**Materials.** All reagents were purchased from commercial vendors and used without further treatment, including triphenylamine, pyridin-4-ylboronic acid, malonic acid, 1-bromopyrrolidine-2,5-dione, tetrakis(triphenylphosphine)palladium(0), 1,4-dioxane methanol, pyridine, piperidine, ethanol, $H_2SO_4$, DCM, DMF.

#### Material synthesis

##### Synthesis of TPACHO

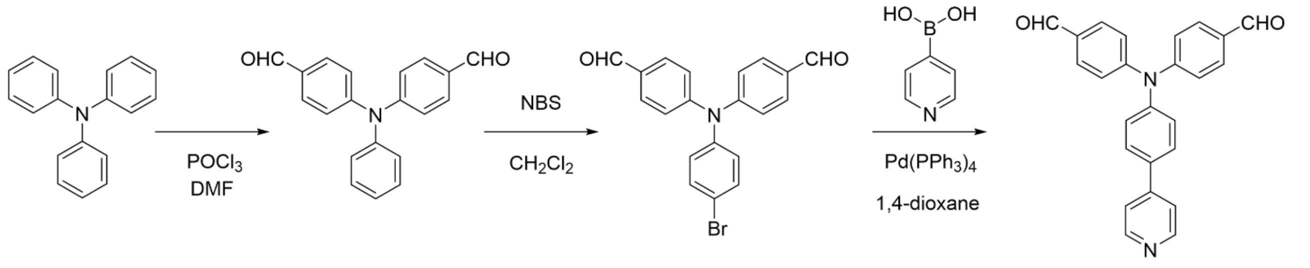

TPACHO

The synthesis of TPACHO followed reported procedures using triphenylamine as the starting material. [1]

##### Synthesis of TPACOOH

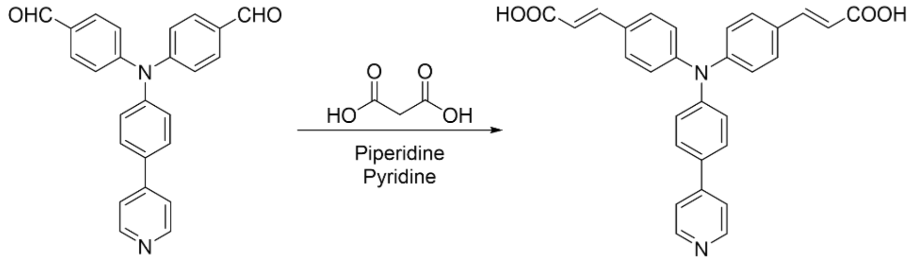

TPACHO TPACOOH

1.13 g (3.0 mmol) TPACHO and malonic acid 0.66 g (6.3 mmol) were dissolved in pyridine (12 mL) and 5 drops of piperidine were added. The above mixture was refluxed and monitored by TLC. After refluxed at 120 ℃ for four days, water was added to the mixture and the pH value of the solution was adjusted to 3.0. Next, the precipitate was purified via recrystallization in ethanol to yield a yellow solid of TPACOOH (0.83 g, yield 60%). $^{1}$H NMR (400 MHz, DMSO) δ 12.33 (s, 2H), 8.73 (d, $J$ = 5.7 Hz, 2H), 7.99 (d, $J$ = 6.1 Hz, 2H), 7.91 (d, $J$ = 8.5 Hz, 2H), 7.68 (d, $J$ = 8.4 Hz, 4H), 7.56 (d, $J$ = 15.9 Hz, 2H), 7.20 (d, $J$ = 8.6 Hz, 2H), 7.10 (d, $J$ = 8.3 Hz, 2H), 6.44 (d, $J$ = 16.0 Hz, 2H). $^{13}$C NMR (100 MHz, DMSO) δ 169.40, 150.19, 147.37, 147.02, 146.18, 139.20, 131.63, 130.65, 129.05, 128.08, 124.18, 123.97, 123.14, 120.63.

##### Synthesis of TPAE

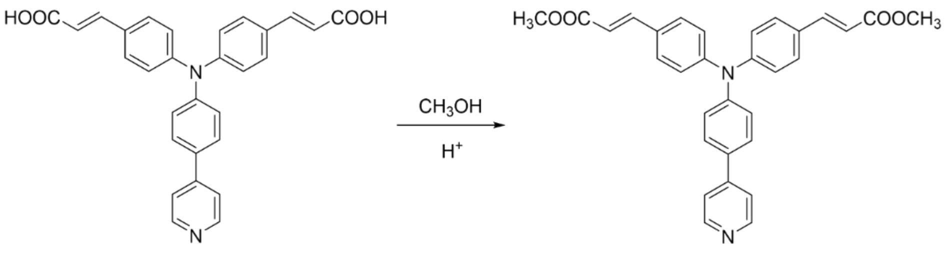

TPACOOH TPAE

253 mg (0.55 mmol) TPACOOH was dissolved in 15 mL methanol. After 15 drops of $H_2SO_4$ were added, the reaction mixture was refluxed at 100 ℃ for 48 h. The reaction solvent was evaporated under vacuum and DCM was added to dissolve the residue. The final product TPAE was obtained after chromatographic purification (115 mg, yield 42%). $^1H$ NMR (400 MHz, $CDCl_3$) δ 8.66 (m, 2H), 7.67 – 7.58 (m, 4H), 7.50 – 7.44 (m, 6H), 7.22 (d, 2H), 7.12 (d, *J* = 8.5 Hz, 4H), 6.35 (d, *J* = 15.9 Hz, 2H), 3.81 (s, 6H). $^{13}C$ NMR (100 MHz, $CDCl_3$) δ167.74, 150.39, 148.64, 147.54, 147.49, 144.11, 133.62, 129.69, 129.57, 128.29, 125.47, 124.19, 121.25, 116.49, 51.83. HR-MS calculated for $(C_{31}H_{26}N_2O_4)^+$ (m/z): 490.197, theoretical: 490.189.

**Physical characterizations.** The $^1H$ NMR and $^{13}C$ NMR spectra were measured on a Bruker Avance III HD Ascend 400 MHz NMR spectrometer. Chemical shifts for protons were referenced to the residual solvent peak ($CDCl_3$, $^1H$ NMR: 7.26 ppm, $D_2O$, $^1H$ NMR: 4.79 ppm, $d_6$-DMSO, $^1H$ NMR: 2.50 ppm), chemical shifts for carbons were referenced to the residual solvent peaks ($CDCl_3$, $^{13}C$ NMR: 77.16 ppm, $d_6$-DMSO, $^{13}C$ NMR: 39.51 ppm). Mass spectrometry was performed on a Bruker Biflex III MALDI-TOFMS instrument. Absorption spectra were collected on an Aglient Cary 8454 spectrophotometer. The emission spectra were measured on a HORIBA Fluorolog QM spectrofluorometer.

**Computational methods**. All calculations were performed with the Gaussian 09W[1] program package employing the DFT method with Becke's three-parameter hybrid functional and Lee-Yang-Parr's gradient corrected correlation functional (B3LYP). [2] 6-31G* basis set was applied for H, C, O and N. The geometries of the singlet ground states of compounds were optimized in $H_2O$ using the conductive polarizable continuum model (CPCM). The local minimum on each potential energy surface was confirmed by frequency analysis. Time-dependent DFT calculations produced the singlet excited states of each compound starting from the optimized geometry of the corresponding singlet ground state, using the CPCM method with $H_2O$ as the solvent. The calculated absorption spectra, electronic transition contributions, and electron density difference maps (EDDMs) were generated by GaussSum 3.0. [3] The electronic orbitals were visualized using VMD 1.9.4a51. [4]

**Cell cultures**. HeLa cells and fibroblasts were cultured in the similar way as. [5] Cells were grown in Dulbecco's modified Eagle's medium (#11965118, DMEM, Thermo Fisher Scientific) supplemented with 10% fetal bovine serum (#26140079, FBS, Thermo Fisher Scientific), penicillin (100 units/ ml), and streptomycin (100 μg/ml; #15140163, 10,000 units/ml, Thermo Fisher Scientific) in a 5% $CO_2$ humidified incubator at 37 ℃.

**Cell treatment and staining**. Cells were seeded on a glass-bottom micro-well dish and incubated with 2 ml of DMEM supplemented with 10% FBS for 24 h, then stained with TPAE (5 μM) for 30 min and with 100 nM MitoTracker Deep Red (MTDR #M7514, Invitrogen) or 200 nM LysoTracker Red (LTR #L7526, Invitrogen) at 37 °C for another 30 min, followed by 10 μM CCCP for 12 h. After treatment, the cells were washed 3 times with pre-warmed free DMEM, and washed with free DMEM 3 times. Finally, cells were cultured in phenol-free medium (#1894117, Gibco) and observed under Nikon-SIM super-resolution microscope (Tokyo, Japan).

**SIM super-resolution microscopy imaging**. Super-resolution images were acquired on a commercial Nikon-SIM Microscope (version AR5.11.00 64 bit, Tokyo, Japan). Images were obtained at 512 × 512 using Z-stacks with a step of 0.2 μm. All fluorescence images were analyzed and their backgrounds were subtracted with ImageJ software.

**Measurement of the distance between lysosomes and mitochondria.** The distances between lysosomes and mitochondria were measured manually by using ImageJ program.

***Supporting figures and tables***

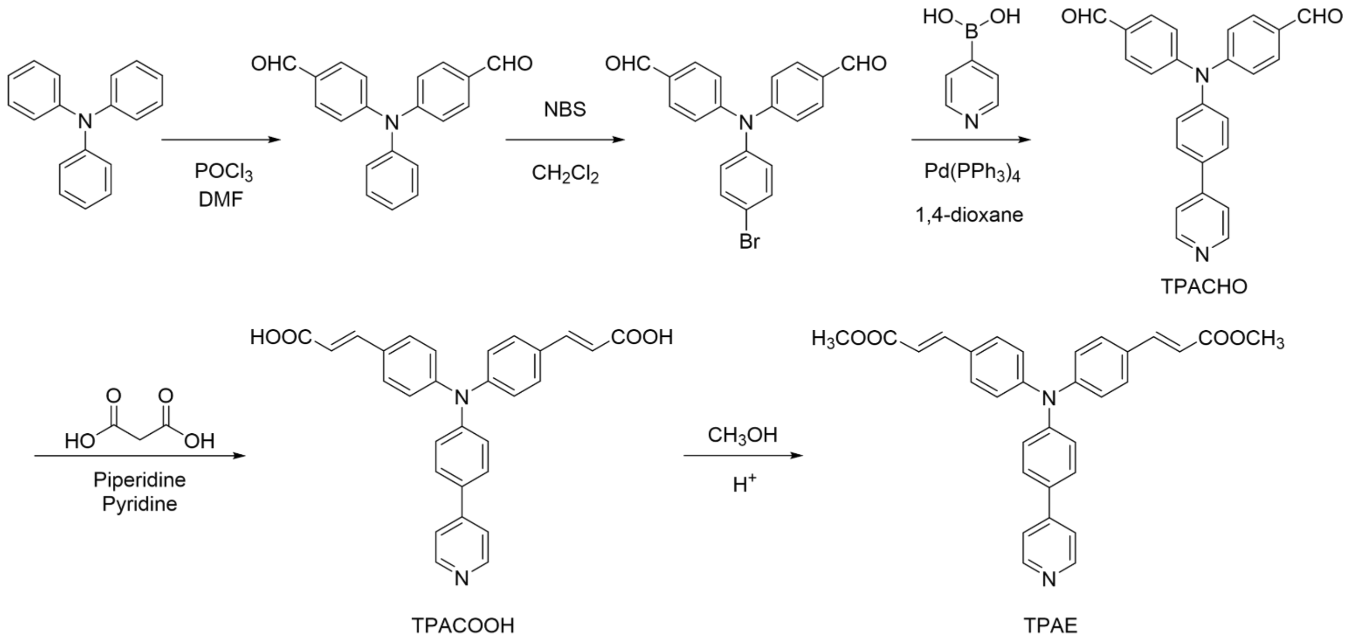

**Figure S1.** The synthetic procedure of TPAE.

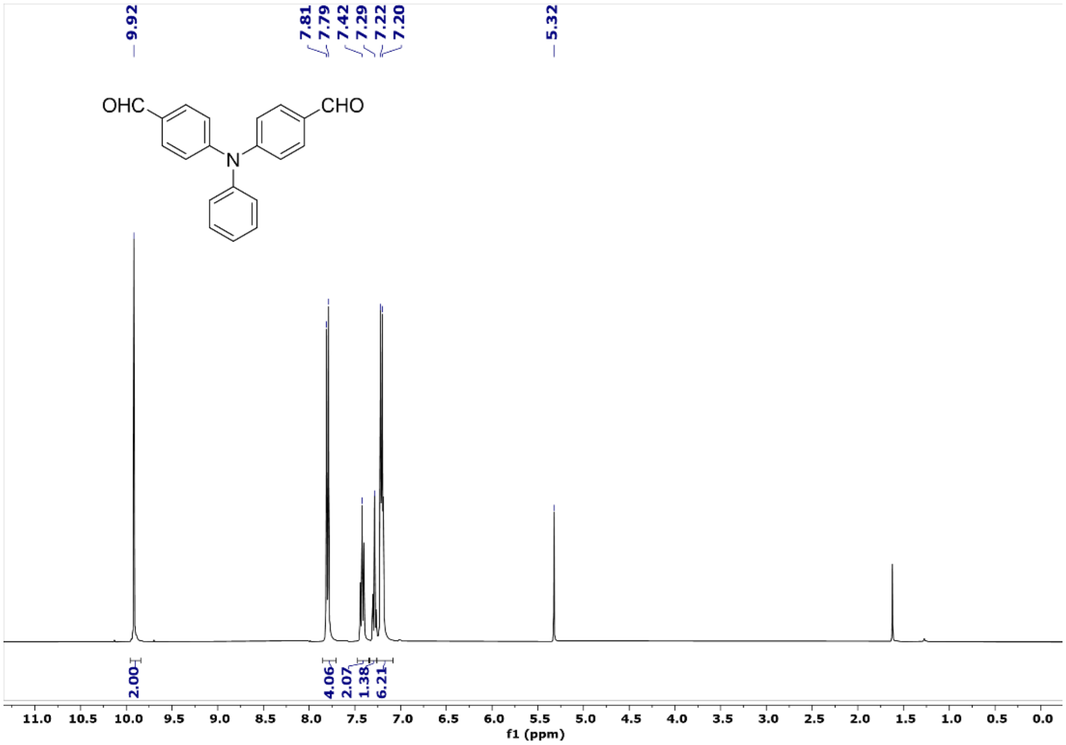

**Figure S2.** $^{1}$H NMR spectrum of 4,4'-(phenylazanediyl)dibenzaldehyde.

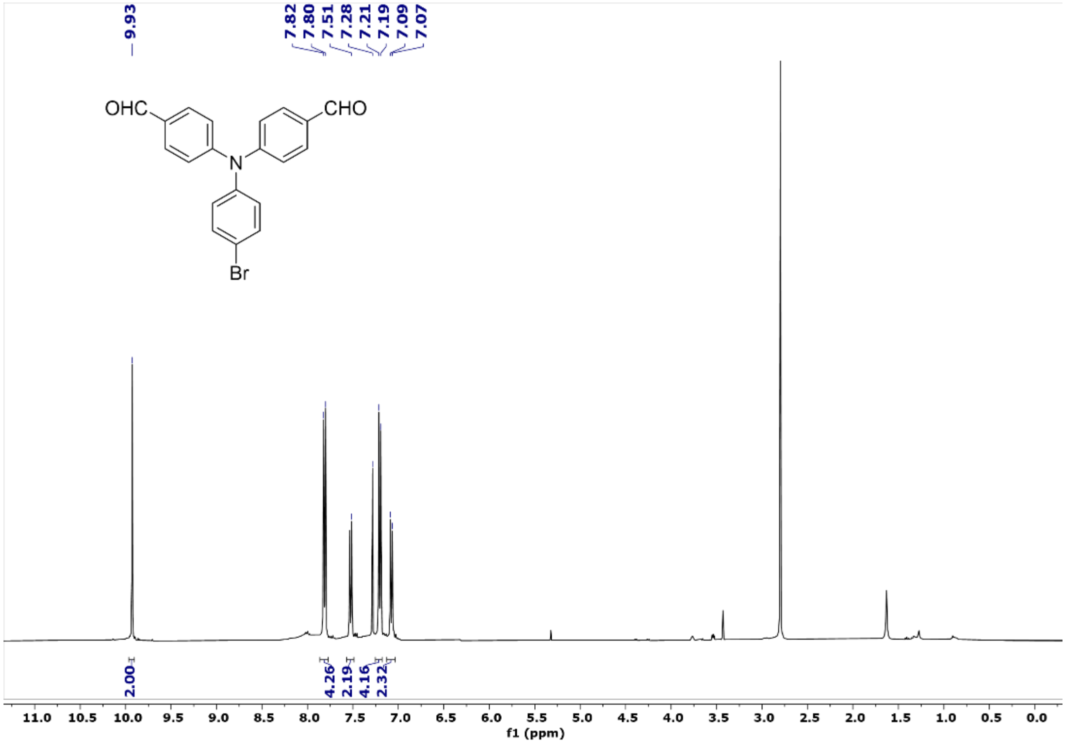

**Figure S3.** $^{1}$H NMR spectrum of 4,4'-((4-bromophenyl)azanediyl)dibenzaldehyde.

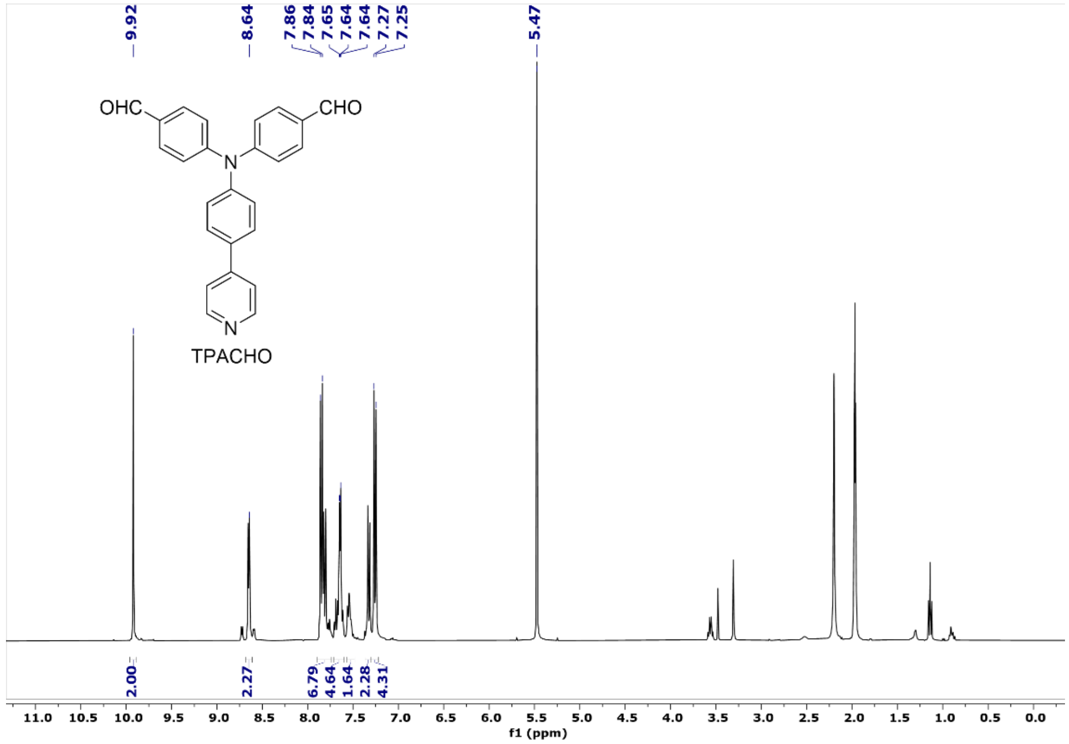

**Figure S4.** $^{1}$H NMR spectrum of TPACHO.

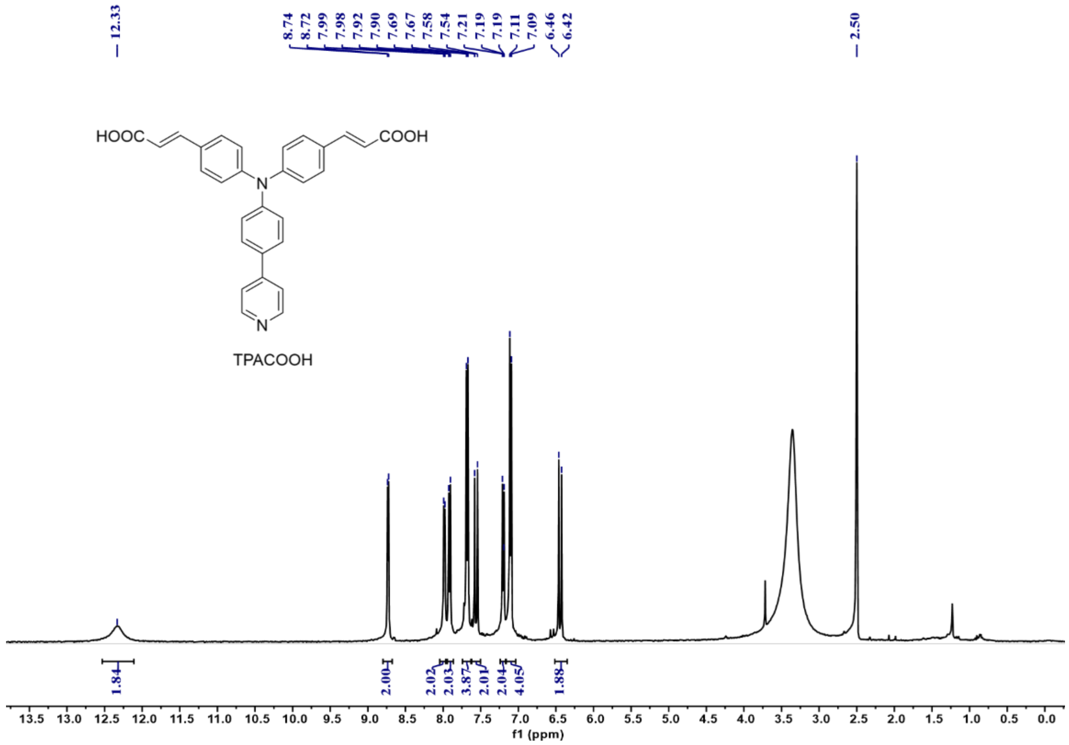

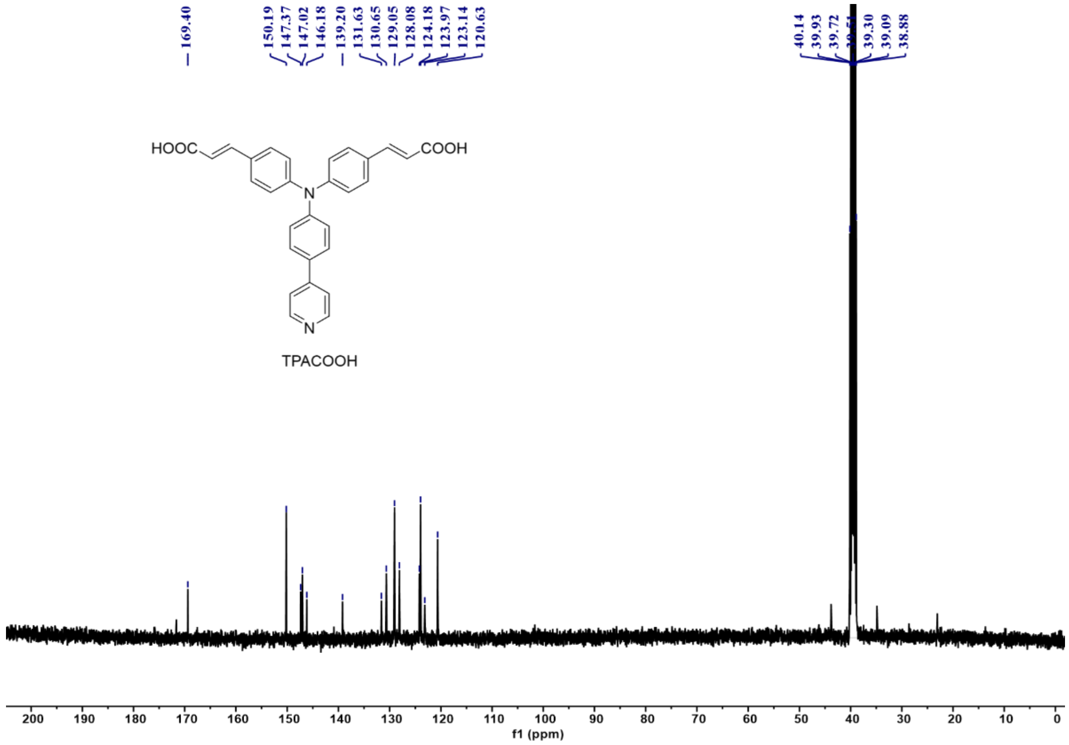

**Figure S5.** $^{1}$H (top) and $^{13}$C (bottom) NMR spectra of TPACOOH.

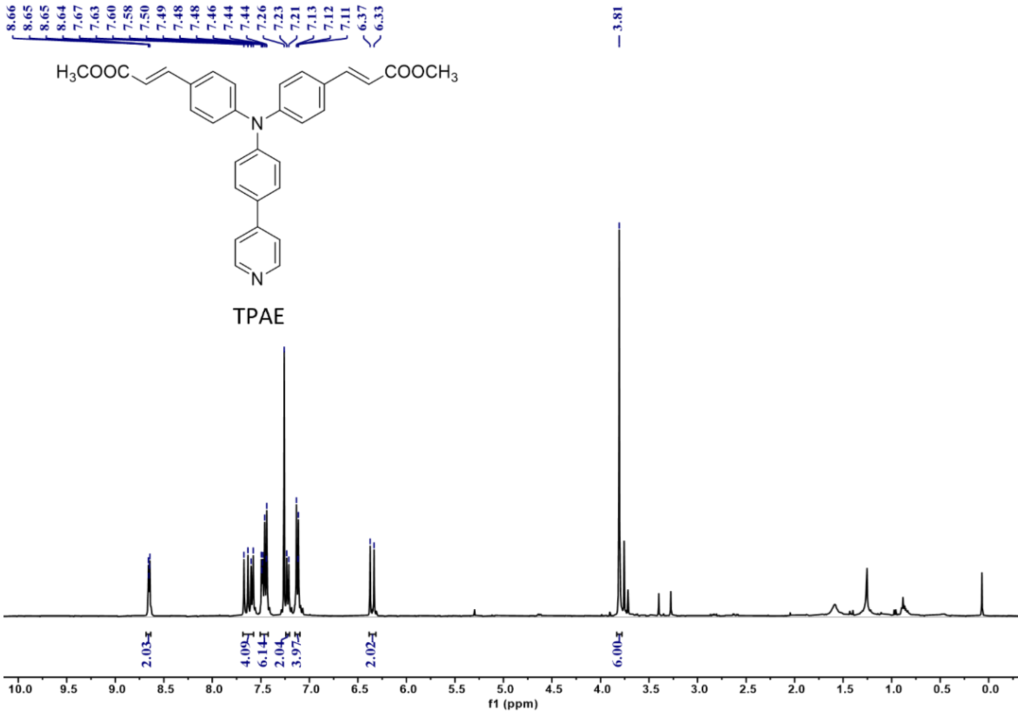

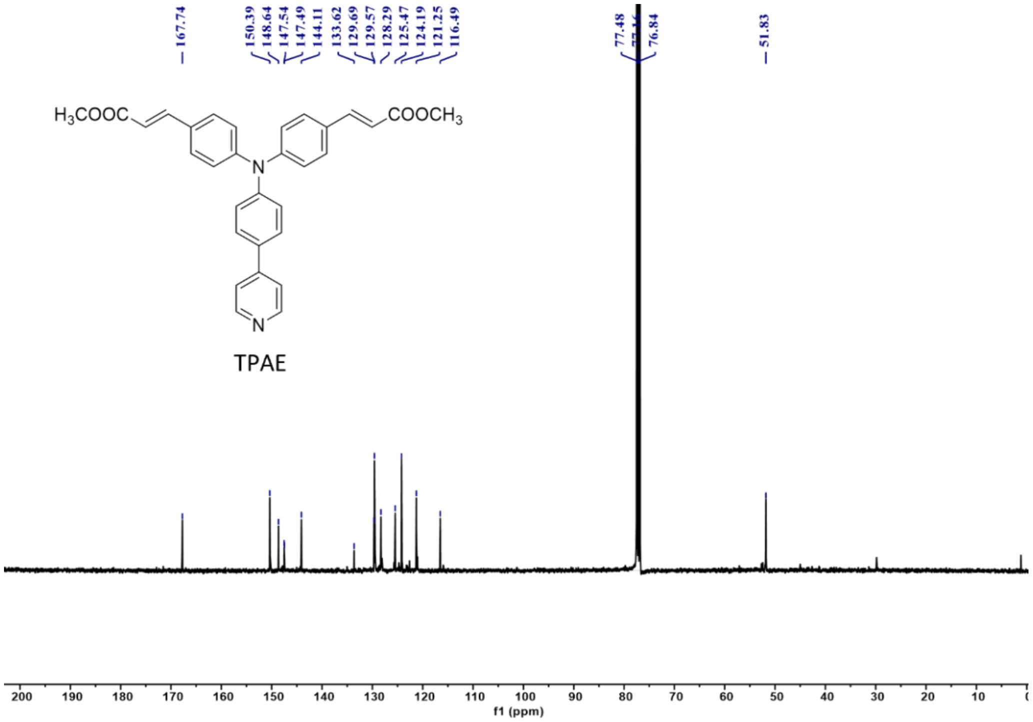

**Figure S6.** $^{1}H$ (top) and $^{13}C$ (bottom) NMR spectra of TPAE.

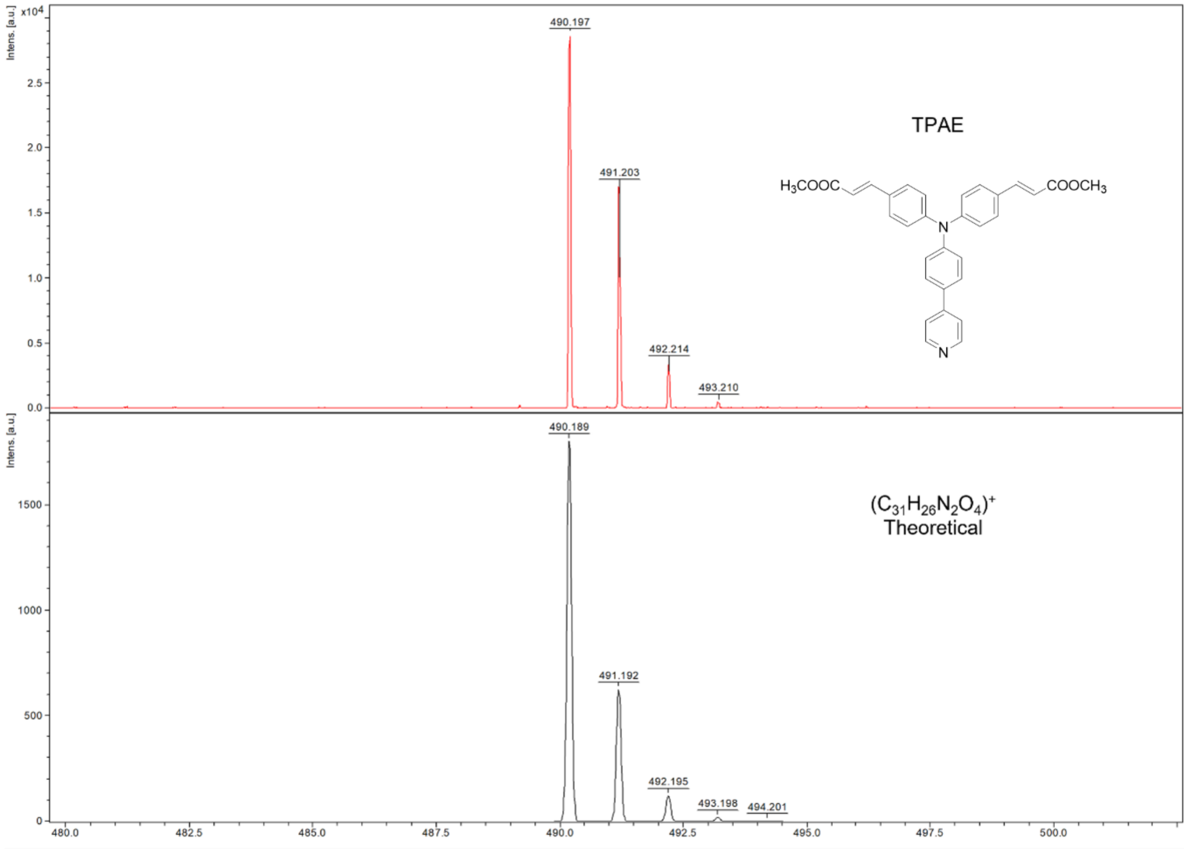

**Figure S7**. Experimental (top, red) and theoretical (bottom, black) HR-MS results of TPAE.

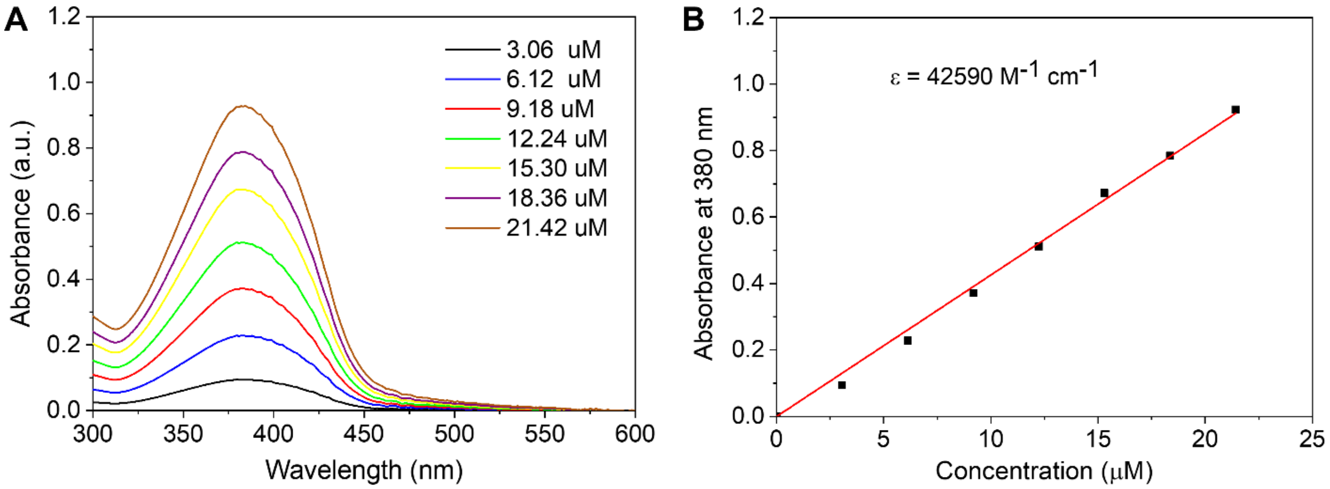

**Figure S8.** (A) Absorption spectra of TPAE with different concentrations in aqueous buffer at pH 7. (B) Linear plot of the maximum absorbance of TPAE at 380 nm against different concentration.

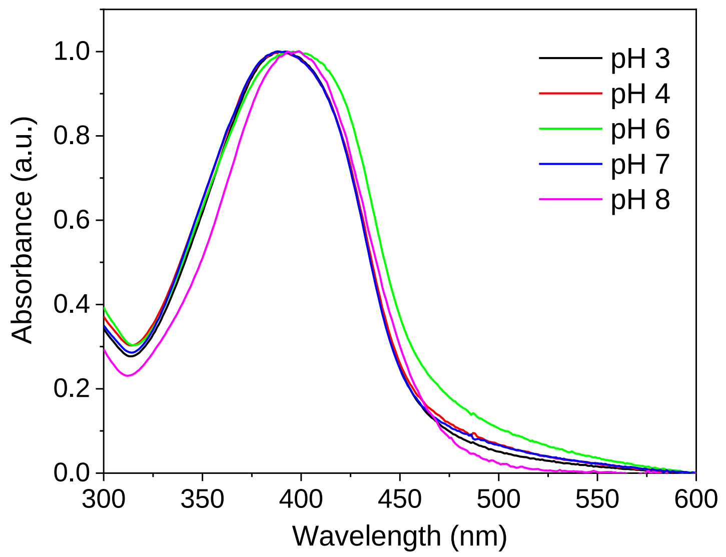

**Figure S9.** Absorption spectra of TPAE in aqueous buffers of different pH.

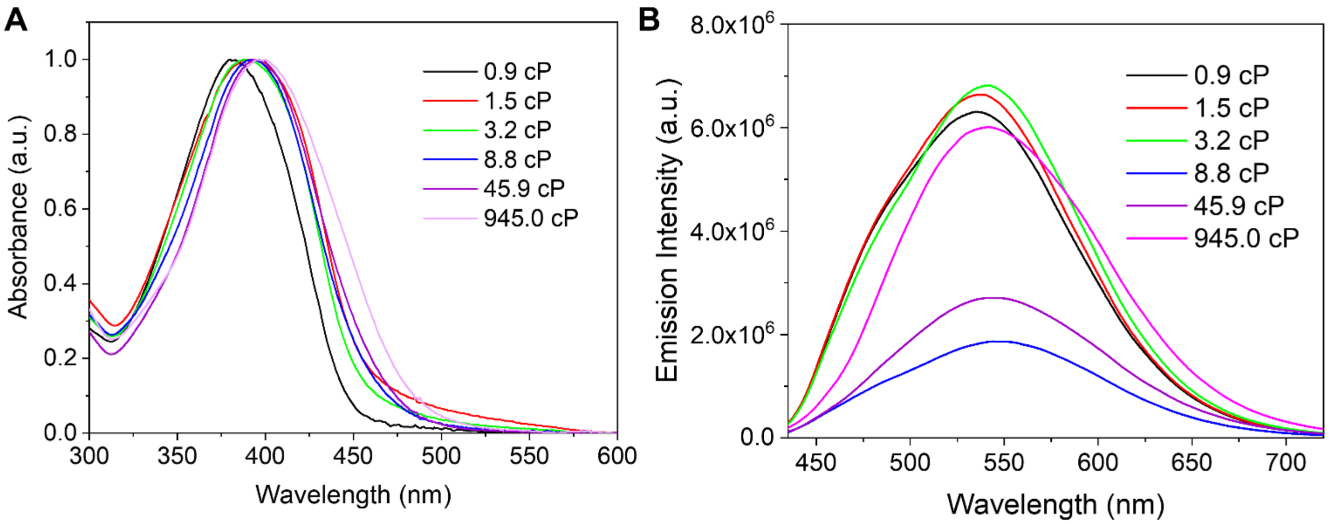

**Figure S10.** Absorption (A) and emission (B) spectra of TPAE (10 μM) in water-glycerol solutions with various glycerol volume percentages.

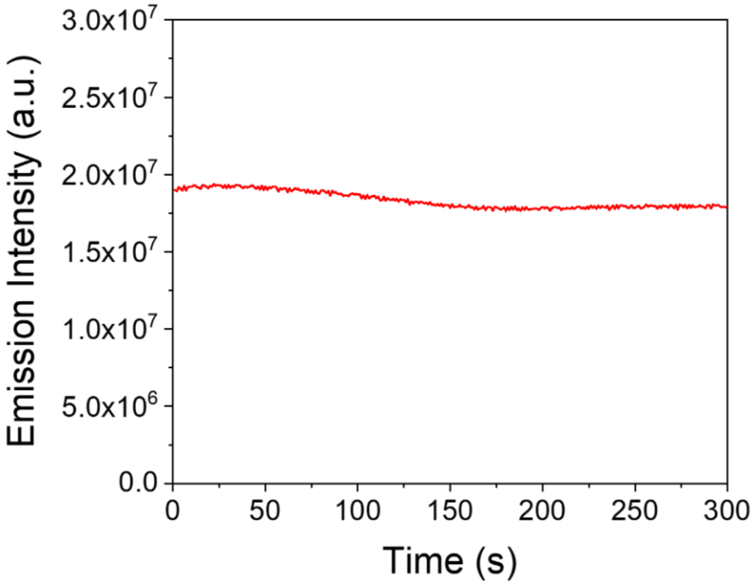

**Figure S11.** The photostability test of TPAE under an irradiation wavelength of 405 nm.

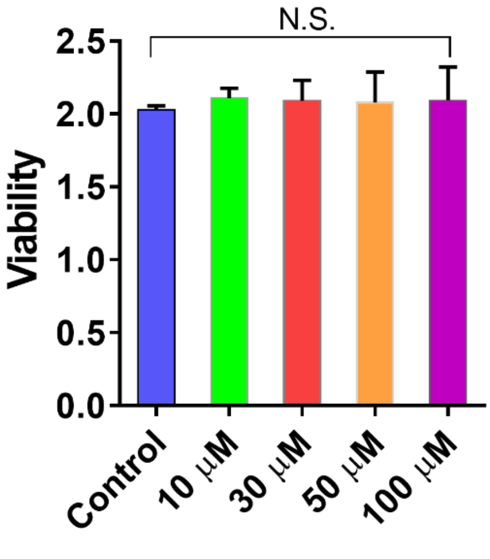

**Figure S12.** Viability of HeLa cells treated with the various concentrations of TPAE for 24 h; measurement was performed with CCK-8 assay (n=6). Data are presented as mean ± SEM; N.S., no significance, unpaired two-tailed t-test.

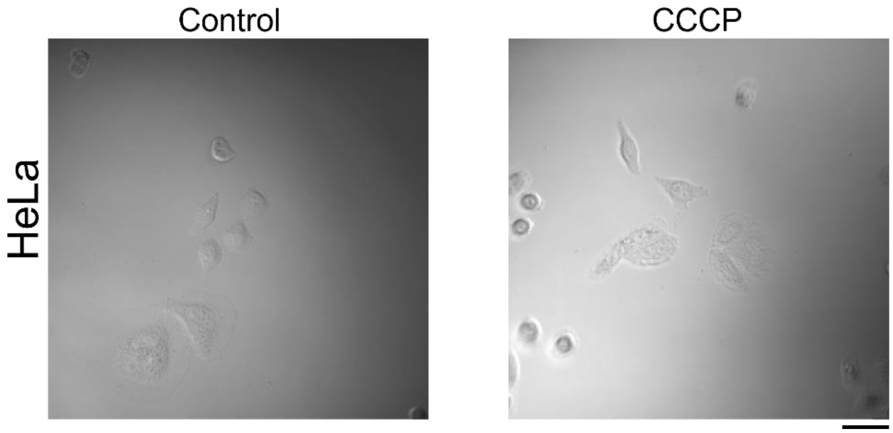

**Figure S13.** Bright filed images of HeLa cells treated with or without CCCP for 12 h. Scale bar, 20 μm.

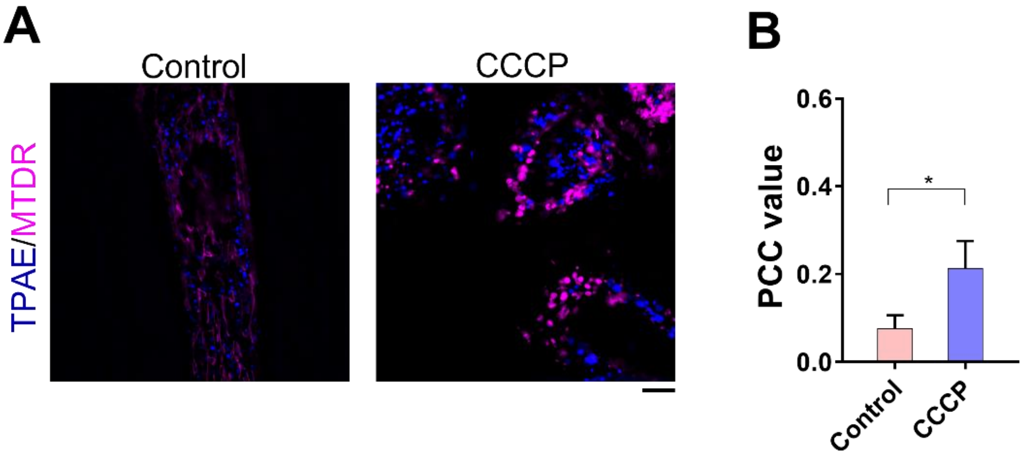

**Figure S14.** (A) SIM imaging of HeLa cells co-stained with TPAE and MTDR after being treated with or without CCCP for 12 h. Scale bar, 5 μm. (B) Pearson correlation coefficient (PCC) value for TPAE and MTDR in HeLa cells. n=6, Data are presented as mean ± SEM. *$p < 0.05$, unpaired two-tailed t-test.

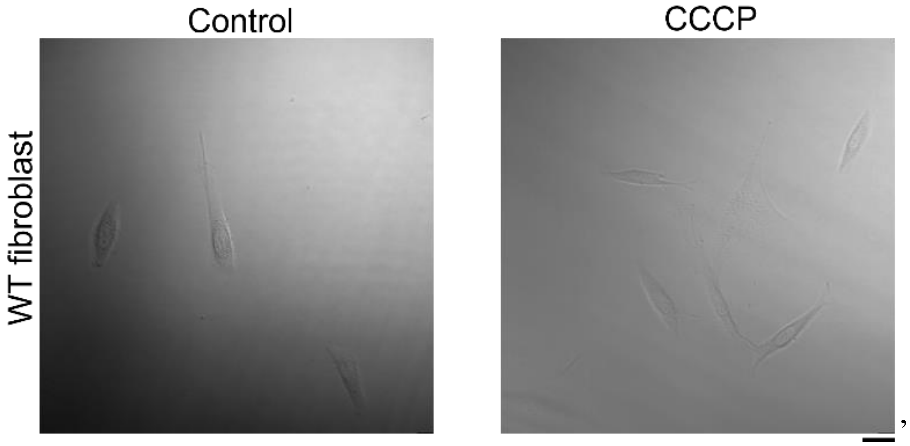

**Figure S15.** Bright filed images of WT human fibroblasts treated with or without CCCP for 12 h. Scale bar, 20 μm.

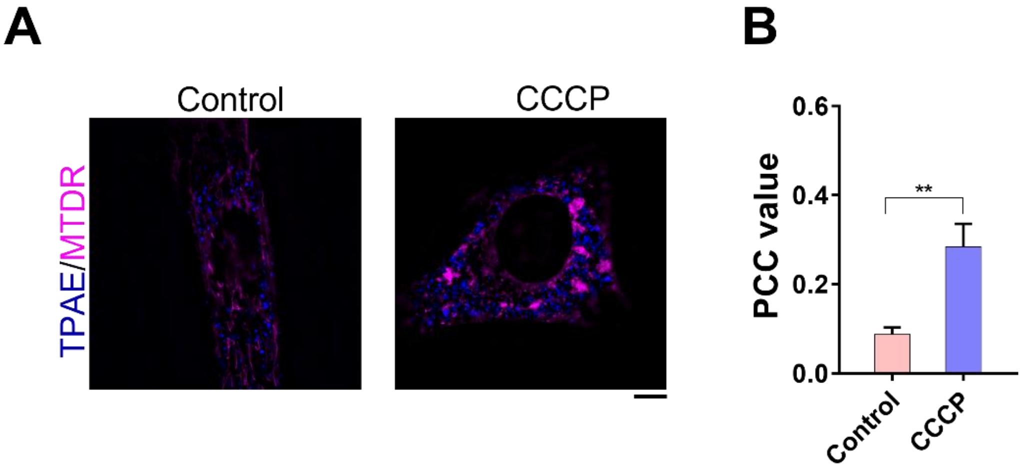

**Figure S16.** (A) SIM imaging of human fibroblasts co-stained with TPAE and MTDR after being treated with or without CCCP for 12 h. Scale bar, 5 μm. (B) PCC value for TPAE and MTDR in human fibroblasts . n=6, Data are presented as mean ± SEM. ***p* < 0.01, unpaired two-tailed t-test.

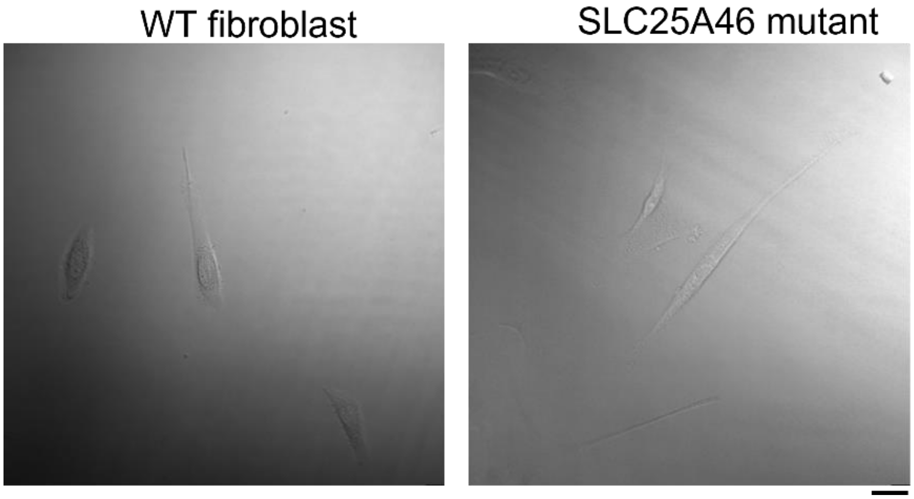

**Figure S17.** Bright filed images of WT human fibroblasts and SLC25A46 mutant fibroblasts. Scale bar, 20 μm.

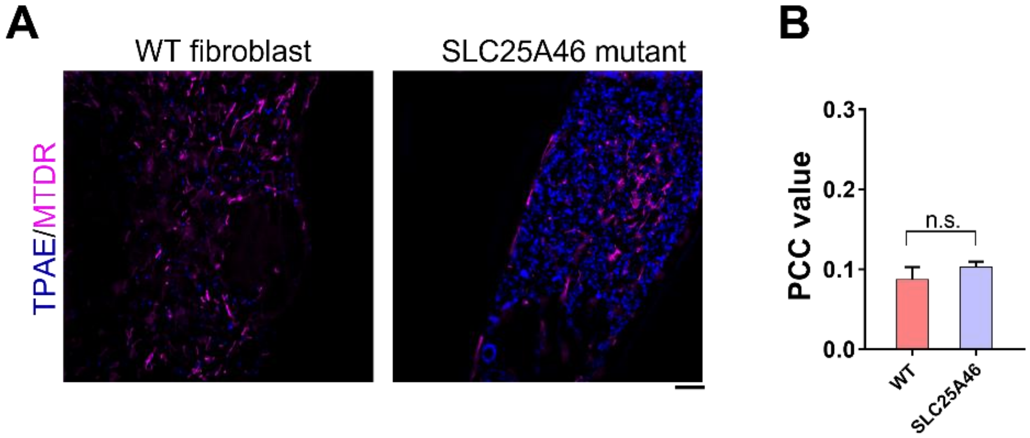

**Figure S18.** (A) SIM imaging of WT fibroblasts and SLC25A46 mutant fibroblasts co-stained with TPAE and MTDR. Scale bar, 5 μm. (B) PCC value for TPAE and MTDR in WT and SLC25A46 mutant fibroblasts, respectively. n=6, Data are presented as mean ± SEM. n.s., no significance, unpaired two-tailed t-test.

**Table S1.** Calculated molecular orbitals of TPAE in $H_2O$.

| MOs | Energy (eV) | Orbitals |
|---|---|---|
| HOMO-5 (124) | -7.172 | 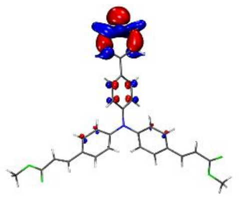 |
| HOMO-4 (125) | -7.147 | 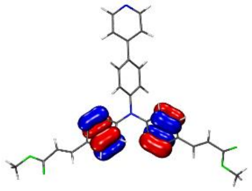 |
| HOMO-3 (126) | -7.089 | 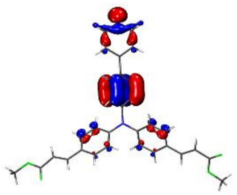 |
| HOMO-2 (127) | -6.720 | 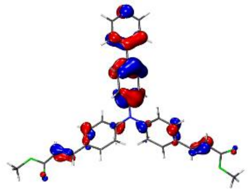 |
| HOMO-1 (128) | -6.567 | 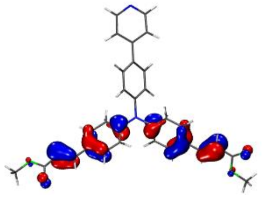 |
| HOMO (129) | -5.275 | 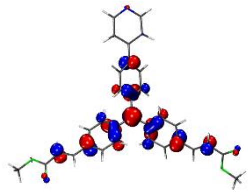 |

| | | |
|---|---|---|
| LUMO (130) | -2.027 | 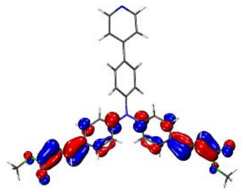 |
| LUMO+1 (131) | -1.798 | 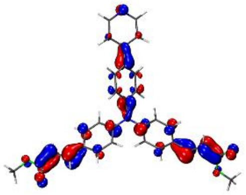 |
| LUMO+2 (132) | -1.242 | 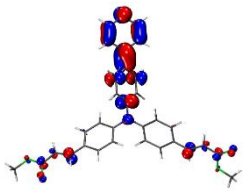 |
| LUMO+3 (133) | -0.706 | 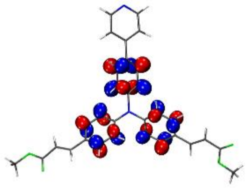 |
| LUMO+4 (134) | -0.377 | 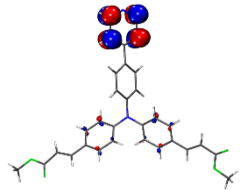 |
| LUMO+5 (135) | -0.139 | 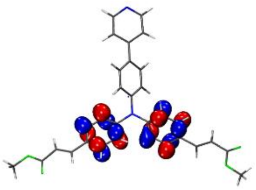 |

**Table S2.** Calculated singlet electron transitions ($f$ > 0.01) of TPAE in $H_2O$.

| No. | Wavelength (nm) | $f$ | Major contributions | EDDM |
|---|---|---|---|---|
| 1 | 437.0 | 1.018 | HOMO->LUMO (99%) | 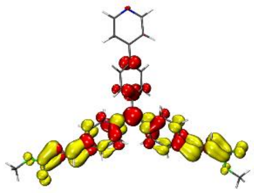 |
| 2 | 404.8 | 0.586 | HOMO->L+1 (97%) | 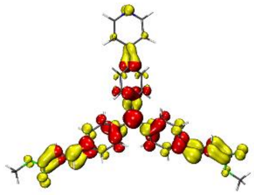 |
| 3 | 338.9 | 0.058 | HOMO->L+2 (93%) | 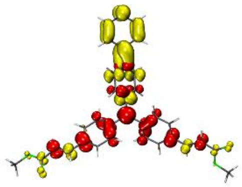 |
| 4 | 320.1 | 0.018 | HOMO->L+3 (92%) | 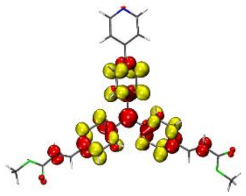 |
| 5 | 292.4 | 0.233 | H-1->LUMO (77%) | 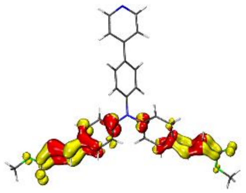 |
| 6 | 290.0 | 0.133 | H-2->LUMO (84%) | 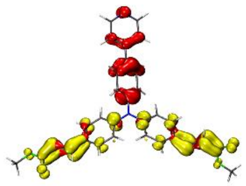 |

| | | | | |
|---|---|---|---|---|
| 7 | 284.7 | 0.024 | H-1->L+1 (17%), HOMO->L+4 (35%), HOMO->L+6 (25%) | 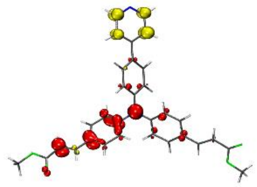 |
| 8 | 284.1 | 0.034 | H-7->LUMO (10%), H-1->LUMO (12%), HOMO->L+5 (52%) | 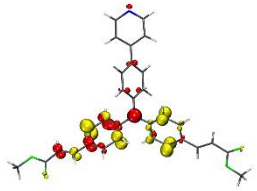 |
| 9 | 282.1 | 0.539 | H-1->L+1 (81%) | 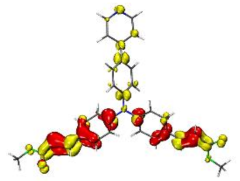 |
| 12 | 274.4 | 0.012 | H-4->LUMO (10%), HOMO->L+4 (49%), HOMO->L+6 (32%) | 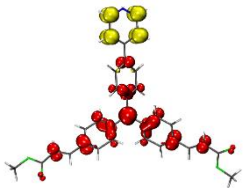 |
| 13 | 270.3 | 0.021 | H-2->L+1 (85%) | 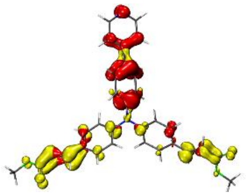 |
| 14 | 267.4 | 0.040 | H-5->L+1 (14%), H-5->L+2 (16%), H-4->LUMO (19%), H-3->L+1 (27%), H-3->L+2 (11%) | 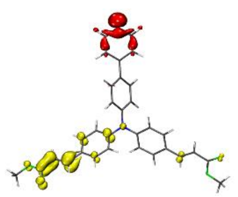 |
| 16 | 265.9 | 0.070 | H-5->L+1 (11%), H-4->LUMO (44%), HOMO->L+6 (21%) | 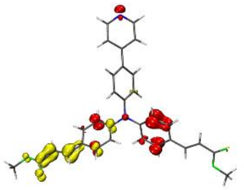 |

| | | | | |
|---|---|---|---|---|
| 17 | 258.3 | 0.018 | H-7->LUMO (29%), H-4->L+1 (21%), HOMO->L+5 (20%) | 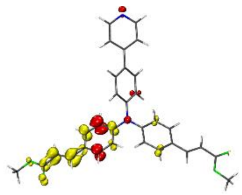 |
| 18 | 256.4 | 0.029 | H-5->L+1 (26%), H-3->L+1 (51%) | 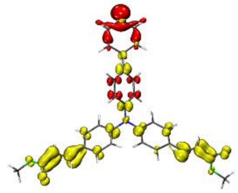 |
| 21 | 251.8 | 0.024 | H-1->L+2 (79%), HOMO->L+8 (13%) | 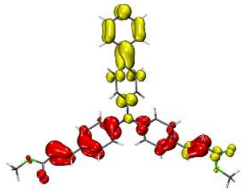 |
| 23 | 249.4 | 0.028 | H-6->L+1 (59%), H-5->L+1 (10%) | 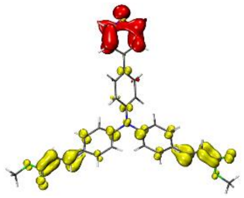 |
| 24 | 249.1 | 0.048 | HOMO->L+7 (87%) | 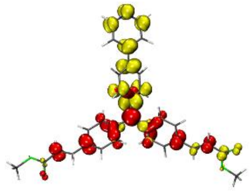 |
| 27 | 245.0 | 0.178 | H-2->L+2 (94%) | 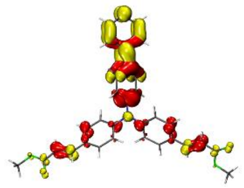 |
| 28 | 239.4 | 0.207 | H-10->LUMO (87%) | 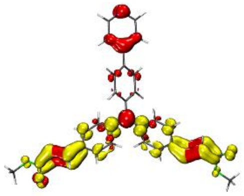 |

**Table S3.** Cartesian coordinates of optimized **TPAE:**

Multiplicity = 1

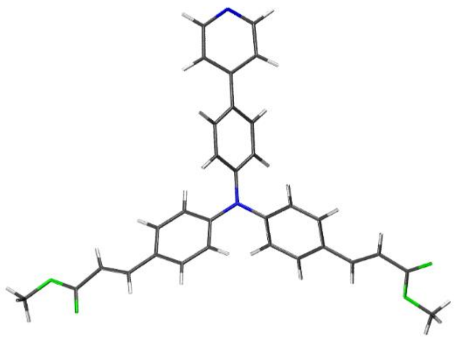

| | | | |
|---|---|---|---|
| N | 0.04259300 | 0.08520900 | 0.04711300 |
| C | -0.02713100 | 1.50696000 | 0.02597100 |
| C | 1.30442600 | -0.55634100 | -0.00058400 |
| C | -1.15086900 | -0.67427400 | 0.11737900 |
| C | -0.88252700 | 2.19244500 | 0.90246100 |
| C | -0.95503900 | 3.58069300 | 0.87177600 |
| C | -0.16658400 | 4.33414700 | -0.01592600 |
| C | 0.69362700 | 3.63599800 | -0.88192900 |
| C | 0.75815900 | 2.24700500 | -0.87149700 |
| C | 1.48700600 | -1.73400700 | -0.74500200 |
| C | 2.72974700 | -2.35321600 | -0.78054700 |
| C | 3.84004600 | -1.82392300 | -0.09419500 |
| C | 3.64394700 | -0.63879800 | 0.64507500 |
| C | 2.40474800 | -0.01969600 | 0.69681100 |
| C | -1.21965300 | -1.83874200 | 0.90086800 |
| C | -2.39655100 | -2.57397800 | 0.95924700 |
| C | -3.55144000 | -2.17778400 | 0.25688600 |
| C | -3.46976900 | -1.00479700 | -0.52230100 |
| C | -2.29657600 | -0.27030500 | -0.59635100 |
| C | 5.11895600 | -2.52016300 | -0.18090900 |
| C | 6.29007100 | -2.15594900 | 0.38403100 |

| | | | |
|---|---|---|---|
| C | 7.54305200 | -2.91398200 | 0.25382600 |
| O | 8.59695200 | -2.56979600 | 0.77064500 |
| O | 7.42534300 | -4.03516500 | -0.49463500 |
| C | 8.62479800 | -4.81188700 | -0.65134300 |
| C | -4.75763300 | -2.98802400 | 0.36731500 |
| C | -5.95905300 | -2.76591900 | -0.20566300 |
| C | -7.07423600 | -3.69968000 | 0.01617900 |
| O | -8.18247800 | -3.29219700 | -0.64670200 |
| O | -7.04257600 | -4.71373900 | 0.69848200 |
| C | -9.34189600 | -4.13040800 | -0.50323400 |
| C | -0.23982600 | 5.81351600 | -0.03802500 |
| C | -1.43348300 | 6.49925900 | 0.24200500 |
| C | -1.44916900 | 7.89162800 | 0.20987700 |
| N | -0.38015300 | 8.64920700 | -0.08034300 |
| C | 0.75884600 | 7.99258800 | -0.34955700 |
| C | 0.88066500 | 6.60511400 | -0.34023500 |
| H | -1.48123100 | 1.63503700 | 1.61543900 |
| H | -1.60634500 | 4.08636800 | 1.57826700 |
| H | 1.29315700 | 4.18230500 | -1.60383300 |
| H | 1.41001200 | 1.73040600 | -1.56843300 |
| H | 0.65573300 | -2.15661200 | -1.29903600 |
| H | 2.85157800 | -3.26027300 | -1.36703100 |
| H | 4.46650800 | -0.20193100 | 1.20286300 |
| H | 2.27684000 | 0.88168200 | 1.28657600 |
| H | -0.35222900 | -2.15985800 | 1.46745400 |
| H | -2.43124000 | -3.46823100 | 1.57609500 |
| H | -4.33051900 | -0.66984800 | -1.09289100 |
| H | -2.25548000 | 0.61920600 | -1.21584100 |
| H | 5.10898800 | -3.43318900 | -0.77241700 |
| H | 6.39060400 | -1.25827800 | 0.98586700 |
| H | 9.40382700 | -4.22255000 | -1.14183600 |

| | | | |
|---|---|---|---|
| H | 8.99043000 | -5.15561400 | 0.31974800 |
| H | 8.34327200 | -5.66143600 | -1.27332400 |
| H | -4.67190400 | -3.87786100 | 0.98973400 |
| H | -6.16081300 | -1.91024200 | -0.84160100 |
| H | -9.64465800 | -4.19176700 | 0.54536200 |
| H | -10.12261200 | -3.65489400 | -1.09670800 |
| H | -9.13785100 | -5.13670600 | -0.87835300 |
| H | -2.34898400 | 5.95878800 | 0.46019300 |
| H | -2.37346500 | 8.42629500 | 0.42251700 |
| H | 1.62613400 | 8.60933600 | -0.57944200 |
| H | 1.84528500 | 6.15124600 | -0.54353400 |
